# Fluorescent pH-sensing bandage for point-of-care wound diagnostics

**DOI:** 10.1101/2023.10.08.561402

**Authors:** Marie-Lynn Al-Hawat, Léo-Paul Tricou, Katia Cherifi, Stéphanie Lamontagne, Minh Tran, Amy Ching Yie Ngu, Gabriela Manrique, Natalie Guirguis, Arturo Israel Machuca-Parra, Simon Matoori

**Author notes:** **Corresponding author** Simon Matoori, Université de Montréal, 2940 Chemin de Polytechnique, Montreal, QC H3T 1J4, #6055.

## Abstract

Diabetic foot ulcers (DFUs) are a serious and prevalent complication of diabetes. Current diagnostic options are limited to macroscopic wound analysis such as wound size, depth, and infection. Molecular diagnostics promise to improve DFU diagnosis, staging, and assessment of treatment response. Here, we developed a rapid and easy-to-use fluorescent pH-sensing bandage for wound diagnostics. In a fluorescent dye screen, we identified pyranine as the lead compound due to its suitable pH-sensing properties in the clinically relevant pH range of 6 to 9. To minimize the release of this dye into the wound bed, we screened a library of ionic microparticles and found a strong adhesion of the anionic dye to a cationic polymeric microparticle. These dye-loaded microparticles showed a strong fluorescence response in the clinically relevant pH range of 6 to 9 and a dye release below 1% after one day in biological media. The dye-loaded microparticles were subsequently encapsulated in a calcium alginate hydrogel to minimize the interaction of the microparticles with the wound tissue. This pH-sensing diagnostic wound dressing was tested on full thickness dorsal wounds of mice, and a linear fluorescence response (R^2^ = 0.9909) to clinically relevant pH values was observed. These findings encourage further development of this pH-sensing system for molecular diagnostics in DFUs.

## 1. Introduction

In diabetes, chronic lower extremity wounds (diabetic foot ulcers, DFUs) are a serious complication with a lifetime incidence of 15-25% among patients with diabetes.^1–4^ The prevalence of DFU was 6% in Medicare patients with diabetes.^5^ With standard wound care (wet gauze, debridement, antibiosis in case of infection), only about half of DFUs heal after seven months and the recurrence rate is high with about 60% after three years.^6,7^ Amputations are required in approximately 20% of patients with moderate or severe DFU, making DFUs the leading cause for non-traumatic lower limb amputations in the United States.^86^ Furthermore, the financial burden of DFU on the health care system is high with Medicare costs for DFU wound care of about 6.2 billion USD per year.^5^

Clinical practice guidelines recommend macroscopic diagnostic evaluations to stage DFUs and assess the need for antibiotic therapy and revascularization.^9^ DFU classification is based on ulcer depth and the implication of other tissues such as tendon, muscle, and bone.^9,10^ Evaluating the presence of infections of the diabetic wound relies on the assessment of non-specific inflammation markers (redness, swelling, warmth) and swaps to determine the microbial composition of the wound. As there is a lack of molecular diagnostics in DFUs, there is high interest in developing new point-of-care diagnostics to evaluate prognosis, disease staging, and evaluation of treatment success as recently reviewed by our group.^11^

Preclinical and clinical studies have highlighted wound pH as a promising emerging therapeutic target in chronic wounds. Clinical studies showed that the pH of chronic wounds is between pH 7 and 9.^12–15^ An alkaline pH was also observed in chronic wounds in diabetic mice.^16^ The basic and slightly alkaline pH found in diabetic wounds promotes the growth of pathogenic bacteria, and antibiotic therapy of wounds lowers wound pH and accelerates wound healing.^12–14,17^ High wound pH reduces wound oxygenation, increases proteolytic enzyme activity, and impairs physiologic macrophage activity within the wound bed.^15^ Recent studies also indicate an inhomogeneous distribution of pH values across the wound with implications for wound healing.^18^

Several potentiometric pH sensors for wound diagnostics were recently developed but their toxic components, the need for a battery, and their limitation to assess pH at a single subregion of the wound limit their clinical usefulness and translation. As there is a growing acceptance of wound pH as a biomarker for wound healing, new diagnostic systems are needed to provide fast wound pH measurements at the point-of-care to enable rapid prognosis and evaluation of treatment response. In this study, we developed a fluorescent pH-sensitive alginate hydrogel for wound pH measurement at the point-of-care. This system is based on pH-sensitive dye-loaded microparticles dispersed in a hydrogel matrix. Once in contact with the wound, wound fluid enters through the hydrogel pores and changes the fluorescence of the dye depending on wound fluid pH. This fluorescence change will be measured using a portable fluorometer for point-of-care pH sensing.

## 2. Materials and Methods

### 2.1. Materials

Pyranine, 5(6)-carboxyfluorescein, N-2-hydroxyethylpiperazine-N-2-ethane sulfonic acid (HEPES), sodium chloride, potassium dihydrogen phosphate, Amberlite IRA-900 MP (styrene-divinylbenzene matrix with benzyltrialkylammonium functionality, *BTA MP*), activated charcoal MP, mesostructured silica MP, zeolite, Amberlite CG50 MP (methacrylic matrix with carboxylic acid functionality, *carboxylate MP*), medium viscosity sodium alginate, and calcium sulfate were purchased from Sigma Aldrich (St. Louis, MO). Sulfo-cyanine 7 carboxylic acid was obtained from Lumiprobe (Cockeysville, MD). IRDye 680RD NHS Ester was obtained from LI-COR Biosciences (Lincoln, NE).

### 2.2. Methods

#### 2.2.1. Screen of fluorescent dye for pH sensitivity

##### Selection of the dye

The absorbance spectrum at different pH values was tested to identify a suitable pH-sensitive dye. Four dyes were tested: pyranine, 5(6)-carboxyfluorescein, IRDye 680RD NHS Ester, and sulfo-cyanine 7 carboxylic acid. Each dye (pyranine 0.4 mM, 5(6)-carboxyfluorescein 0.05 mM, IRDye 680RD NHS Ester 0.02 mM, sulfo-cyanine7 carboxylic acid 0.02 mM) was incubated at pH 6 to 9 in isotonic sodium chloride containing phosphate buffer 150 mM (pH 6 – 8) and isotonic sodium chloride containing HEPES buffer 150 mM (pH 8.5, 9). Absorbance spectra were recorded between 400 nm and 900 nm using a plate reader (Spark Multimode Microplate Reader, Tecan, Männedorf, Switzerland).

#### 2.2.2. Screen of microparticles for dye adsorption

To select the most suitable microparticulate dye carrier, the loading of pyranine was assessed for BTA microparticle (MP), activated charcoal MP, mesostructured silica MP, zeolite MP, carboxylate MP. Each microparticle was incubated in 7.5 µM of pyranine at 37°C for 15 min and protected from light. The dispersion was then purified from unbound dye by centrifugation (5 times at 15 000 x g). The dye concentration in the supernatant was quantified by fluorescence and used to calculate pyranine adsorption. Fluorescence was measured using excitation wavelengths of 413 nm (isosbestic) and 455 nm (pH-dependent) and an emission wavelength of 510 nm using a plate reader (Spark Multimode Microplate Reader, Tecan, Männedorf, Switzerland).

#### 2.2.3. pH sensitivity and stability of pyranine-loaded BTA microparticles

To determine the pH sensitivity, we measured the fluorescence of pyranine and pyranine-loaded MP at pH 6 to 9 in isotonic sodium chloride containing phosphate buffer 150 mM (pH 6 – 8) and isotonic sodium chloride containing HEPES buffer 150 mM (pH 8.5, 9). Pyranine-loaded MP were additionally investigated in the presence of 10% FBS at the same pH values after incubation at 37°C for 15 min. Fluorescence was measured using excitation wavelengths of 413 nm (isosbestic) and 455 nm (pH-dependent) and an emission wavelength of 510 nm using a plate reader.

#### 2.2.4. pH sensitivity and stability of pyranine-loaded BTA microparticles loaded into alginate hydrogels

Ionically crosslinked calcium hydrogels were prepared by mixing 120 µL of medium viscosity sodium alginate 8% in water and 40 µL of a calcium sulfate dihydrate suspension 192 mM in water with final concentrations of 6% alginate and 48 mM calcium sulfate and incubation in a cylindrical container at 37°C for 24 hours. Subsequently, the hydrogel was incubated in an isotonic calcium chloride solution 25 mM for 24 hours at 37°C. The pH sensitivity of the pyranine-loaded BTA microparticles-encapsulating hydrogel was determined at pH 6 to 9 with and without FBS 10% (v/v) as explained above.

#### 2.2.5. Investigation of pH sensing in mouse wounds *ex vivo*

Excisional full thickness mouse wounds were performed using a biopsy punch with a diameter of 10 mm. Hydrogels were incubated in isotonic sodium chloride-containing phosphate buffer 150 mM at pH 6, 7, and 8, and in isotonic sodium chloride-containing HEPES buffer 150 mM at pH 9. After 5 min, the hydrogels were placed on the wounds. An image was taken using the LabeoTech OiS300 In Vivo Imaging System (Labeo Technologies Inc, Montreal, QC) device using an excitation wavelength of 466 nm (40 nm bandwidth) and an emission wavelength of 525 nm (50 nm bandwidth).

#### 2.2.6 Statistical analysis

GraphPad and Microsoft Excel 2016 (linear regression analysis) were used for the statistical analysis. Comparisons of three or more groups were performed by a one-way ANOVA followed by Tukey’s post-hoc test. Two groups were compared using a two-sided t test. To compare fluorescence curves, relative fluorescence intensities were compared at pH 7.5. A p-value of <0.05 was deemed statistically significant.

## 3. Results and Discussion

### 3.1. Screen of fluorescent dye for pH sensitivity

To identify a pH-sensitive dye with a fluorescence response between pH 6 and 9, we screened a small library of commercial pH-sensitive dyes. The investigated cyanine dyes sulfo-cyanine 7 and IRDye 680RD were not responsive to pH changes in the targeted range (Figure 1AB) which is likely due to missing electron donors. The fluorescent dye carboxyfluorescein showed a pH-sensitive fluorescence in accordance with the literature^19^ (Figure 1C). However, the pH sensitivity was in a pH range of 6 to 7 and therefore too acidic for our purposes. The pH sensitivity of pyranine was in a pH range of 6 to 9 in accordance with the literature^20^ and thus deemed suitable for our system (Figure 1D, Supplementary Figure S1). An additional advantage of pyranine is its isosbestic excitation wavelength which allows for correcting for small differences in dye concentration. We therefore selected pyranine as the lead compound for our study.

**Figure 1.**
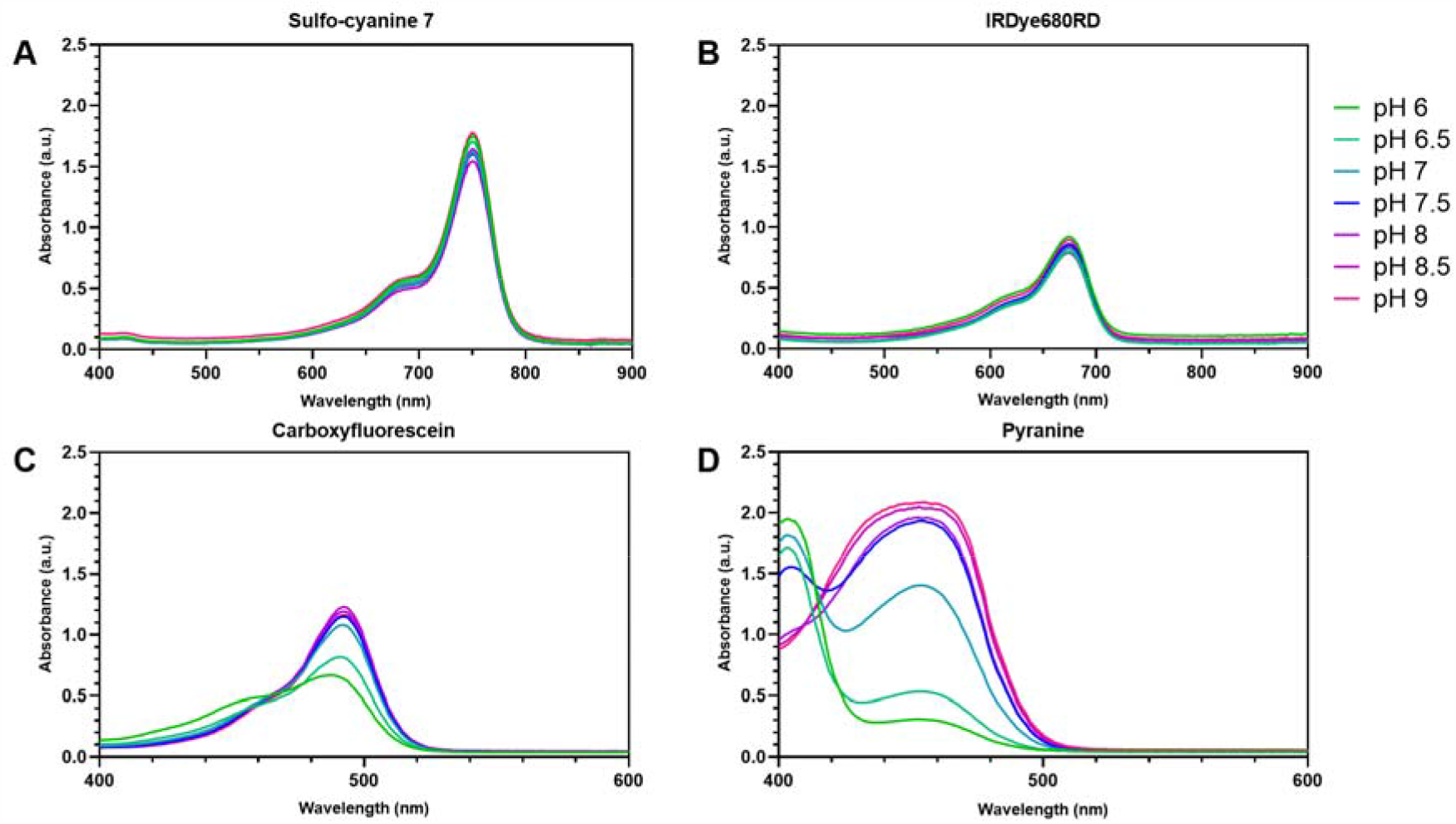
Screen of fluorescent dyes. Absorbance spectrum of sulfo-cyanine 7 (A), IRDye680RD (B), carboxyfluorescein (C) and pyranine (D) at different pH values.

### 3.2. Screen of microparticles for dye adsorption

To identify a carrier with low release of the dye, we screened a small library of ionic microparticles with diameters in the micrometer range (Supplementary Table S1). We saw low loading capacities for zeolites (Figure 2). While zeolites were shown to adsorb anionic molecules, they were limited to small ions such as hydroxide^21^. We hypothesize that pyranine was too large to be efficiently incorporated into the zeolite lattice. Furthermore, there was an ionic repulsion between the anionic pyranine and the anionic surface of the zeolites. Similarly, pyranine was not efficiently adsorbed onto the negatively charged surface of silica and polycarboxylate microparticles. A strong adsorption of pyranine was observed with activated charcoal as expected for this material which is a clinically used adsorbent for charged organic molecules such as drugs^20,22^. However, no fluorescence of the activated charcoal particles was observed after loading with pyranine (Supplementary Figure S2). We hypothesize that the mesoporous structure of activated charcoal leads to a strong loading in inner pores that are not accessible for excitation photons. We observed a good adsorption of pyranine onto cationic BTA microparticles. As pyranine-loaded BTA microparticles exhibited a strong fluorescence (Supplementary Figure S2), we selected these cationic microparticles for our study.

**Figure 2.**
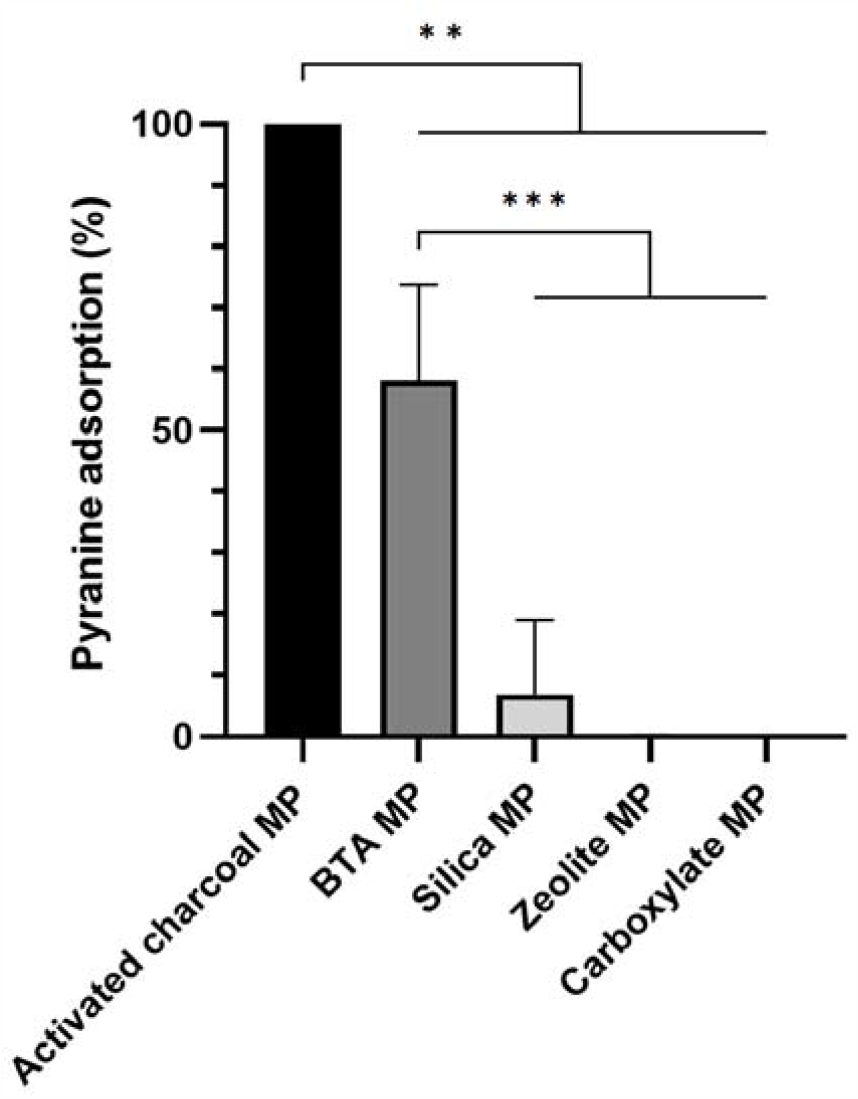
Screen of microparticles for pyranine adsorption. Pyranine adsorption on activated charcoal, BTA, silica, zeolite, and carboxylate microparticles. All results as means ± SD (n = 3). **p < 0.01; ***p < 0.001.

### 3.3. pH sensitivity and stability of pyranine-loaded BTA microparticles

To investigate the pH sensitivity of pyranine-loaded BTA microparticles, they were incubated in buffer solutions from pH 6 to 9. A strong fluorescence response to the pH increase was observed in this pH range (Figure 3A). The response in biological media was also investigated. After the addition of 10% FBS, a similar response curve was observed (Figure 3B). As a release of pyranine into the wound is not desired because dye release may lead to a loss of function, the release of pyranine from the microparticles was investigated in a biological fluid. Very low dye release was observed in the presence of 10% and 50% FBS (Figure 3B). These data indicate that the dye release is likely very low even in the presence of ions and proteins in wound fluid.

**Figure 3.**
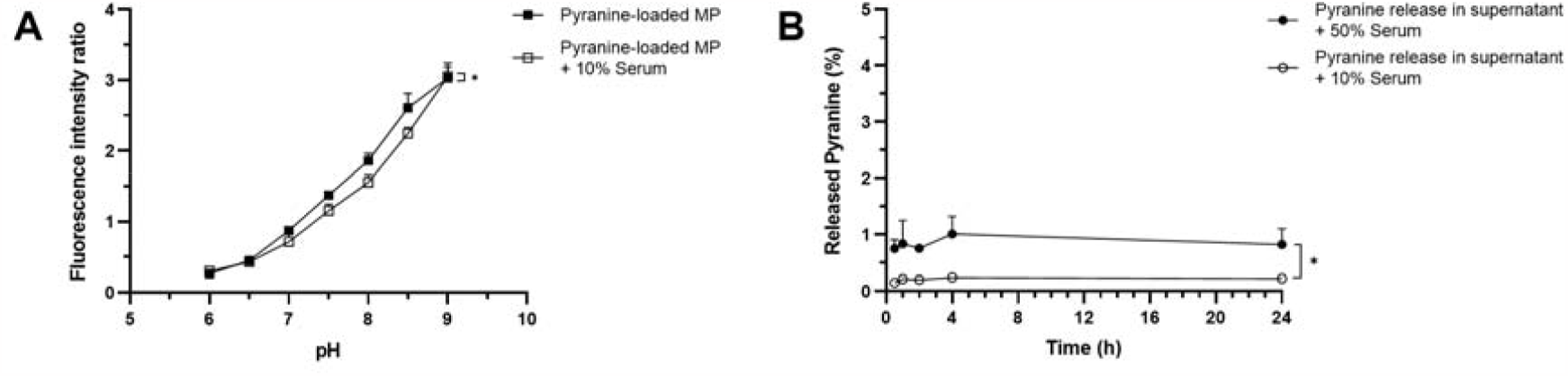
pH sensitivity and stability of pyranine-loaded BTA microparticles. Fluorescence intensity ratio of pyranine-loaded BTA microparticles at different pH values in PBS and in FBS-containing PBS (A). Percentage of released pyranine at different time points after incubation in FBS-containing PBS at 37C (B). All results as means ± SD (n = 3). *p < 0.05.

### 3.4. pH sensitivity and stability of pyranine-loaded BTA microparticles loaded into alginate hydrogels

To reduce their interactions with the wound bed, the microparticles were encapsulated in a calcium-crosslinked alginate hydrogel (Figure 4A). This hydrogel was selected due to its nanoporous structure that retains the microparticles in the hydrogel scaffold. Moreover, the calcium alginate hydrogel was selected for its high biocompatibility and low immunogenicity.^23^ Indeed, a calcium-crosslinked alginate hydrogel is approved by the Food and Drug Administration as a wound dressing for DFU.^24^ The pyranine-loaded BTA microparticles encapsulated in the calcium-crosslinked alginate hydrogel showed a pH-responsive fluorescence between pH 6 and 9 in PBS and in the presence of FBS (Figure 4B). The kinetics of pH sensing was fast with an equilibrium reached after 1 min (Supplementary Figure S2). These data encouraged a further evaluation on mouse wounds.

**Figure 4.**
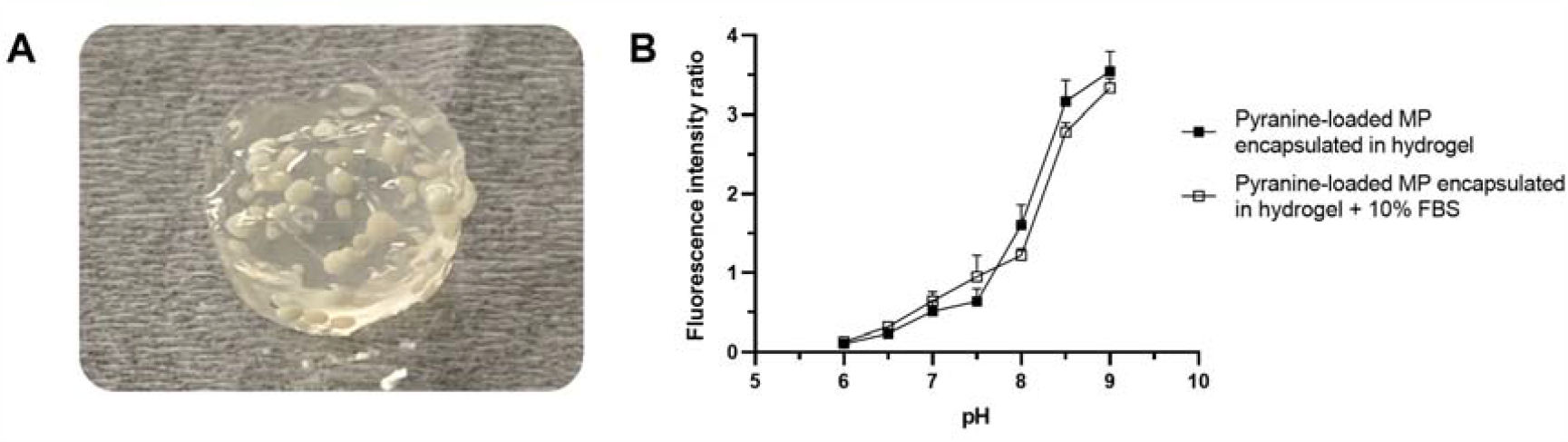
pH sensitivity of pyranine-loaded BTA microparticles-encapsulating calcium alginate hydrogel. Photograph of pyranine-loaded BTA microparticles-encapsulating calcium alginate hydrogel (A). Fluorescence intensity ratio of pyranine-loaded BTA microparticles encapsulating calcium alginate hydrogel at different pH values in PBS and in FBS-containing PBS (B). All results as means ± SD (n = 3).

### 3.5. Investigation of pH sensing in mouse wounds *ex vivo*

After developing a pH-sensitive hydrogel based on pyranine-loaded microparticles encapsulated in a calcium alginate hydrogel, we tested its pH-sensing capacity upon topical administration. We used a full-thickness dorsal wound model in mice. This model is widely established for wound healing studies.^25,26^ Using a small animal fluorescence imaging device, a strong fluorescence response of the pH-sensitive hydrogel with increasing pH was observed (Figure 5A). Quantifying the signal intensity showed a high linearity of the signal in the clinically relevant pH range of 6 to 9 (Figure 5B). The pyranine-loaded microparticles were visible in the fluorescence image. This finding indicates that this diagnostic hydrogel may be capable of visualizing the pH distribution across the wound. In the light of recent studies indicating that the pH of chronic wounds is not homogeneous, the distribution of wound pH may provide valuable additional diagnostic information.

**Figure 5.**
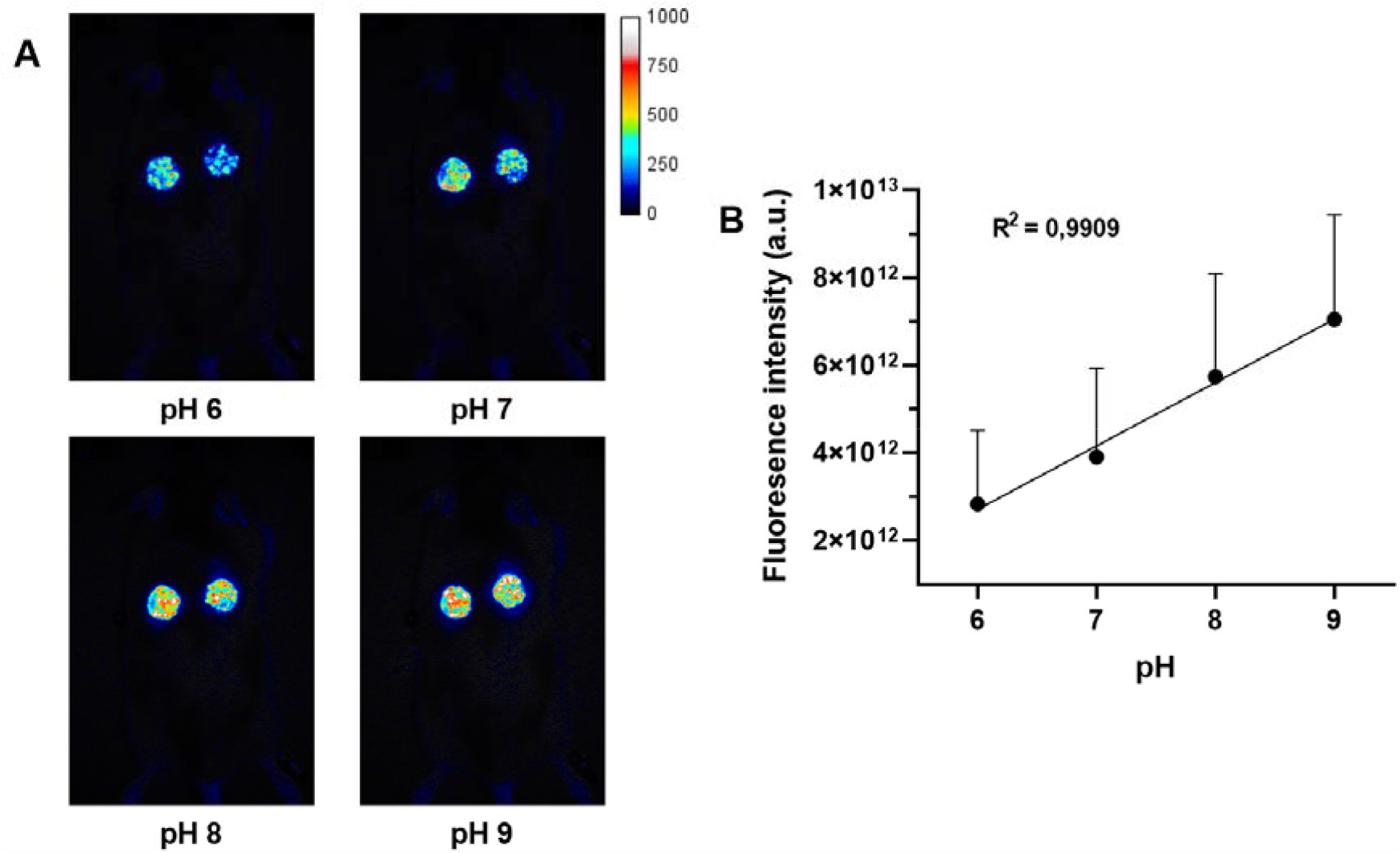
Investigation of pyranine-loaded BTA microparticles-encapsulating calcium alginate hydrogel in mouse wounds *ex vivo*. Fluorescence images of pyranine-loaded BTA microparticles-encapsulating calcium alginate hydrogel placed on dorsal full thickness excisional wounds on C57/BL6 mouse (A). Quantification of fluorescence intensity of pyranine-loaded BTA microparticles encapsulating calcium alginate hydrogel at different pH values (B). All results as means ± SD (n = 5).

## Conclusions

DFUs are a serious and prevalent complication of diabetes. Current diagnostic options for foot ulcers are limited to macroscopic wound analysis. Molecular diagnostics promise to improve DFU diagnosis, staging, and assessment of treatment response. We developed a rapid and easy-to-use pH-sensing bandage for wound diagnostics. This system relies on the loading of a pH-sensitive fluorescent dye onto ionic microparticles. These dye-loaded microparticles showed a strong fluorescence response in the clinically relevant pH range of 6 to 9. Furthermore, the dye release from microparticles was below the targeted dye release of 1% after one day in biological media. Dye release is not desired because leakage impairs the performance of the diagnostic system and exposes the wound to the dye. The dye-loaded microparticles were subsequently encapsulated in a calcium alginate hydrogel to minimize the interaction of the microparticles with the wound tissue. This pH-sensing diagnostic wound dressing was tested on full thickness dorsal wounds of mice, and a linear fluorescence response to clinically relevant pH values was observed. These findings encourage further development of this pH-sensing system for molecular diagnostics in DFUs.

## Acknowledgements

We thank Dr Marc Saba El Leil for the support in the ex vivo study. SM gratefully acknowledges funding from Natural Sciences and Engineering Research Council of Canada (Discovery Grant RGPIN-2022-04384), Canada Foundation for Innovation (Fonds des leaders John-R.-Evans 42712), Fonds de recherche du Québec – Nature et technologies (Relève professorale 330153). NG gratefully acknowledges master’s scholarships from the Groupe de recherche universitaire sur le médicament at Université de Montréal, the Faculté de Pharmacie at Université de Montréal, Canadian Institutes of Health Research, and Fonds de recherche du Québec – Santé.

## References

(1) Baltzis, D.; Eleftheriadou, I.; Veves, A. Pathogenesis and Treatment of Impaired Wound Healing in Diabetes Mellitus: New Insights. Adv. Ther. 2014, 31 (8), 817–836. 10.1007/s12325-014-0140-x.

(2) Matoori, S.; Veves, A.; Mooney, D. J. Advanced Bandages for Diabetic Wound Healing. Sci. Transl. Med. 2021, 13 (585), eabe4839. 10.1126/scitranslmed.abe4839.

(3) Matoori, S. Diabetes and Its Complications. ACS Pharmacol. Transl. Sci. 2022, 5 (8), 513–515.10.1021/ACSPTSCI.2C00122/ASSET/IMAGES/LARGE/PT2C00122_0002.JPEG.

(4) Freedman, B. R.; Hwang, C.; Talbot, S.; Hibler, B.; Matoori, S.; Mooney, D. J. Breakthrough Treatments for Accelerated Wound Healing. Sci. Adv. 2023, 9 (20), eade7007. 10.1126/SCIADV.ADE7007/ASSET/5DB7B785-0E2C-466B-AC37-8DB2D108D9A6/ASSETS/IMAGES/LARGE/SCIADV.ADE7007-F2.JPG.

(5) Nussbaum, S. R.; Carter, M. J.; Fife, C. E.; DaVanzo, J.; Haught, R.; Nusgart, M.; Cartwright, D. An Economic Evaluation of the Impact, Cost, and Medicare Policy Implications of Chronic Nonhealing Wounds. Value Heal. 2018, 21 (1), 27–32. 10.1016/j.jval.2017.07.007.

(6) Armstrong, D. G.; Boulton, A. J. M.; Bus, S. A. Diabetic Foot Ulcers and Their Recurrence. N. Engl. J. Med. 2017, 376 (24), 2367–2375. 10.1056/NEJMra1615439.

(7) Tchanque□Fossuo, C. N.; Dahle, S. E.; Lev□Tov, H.; West, K. I. M.; Li, C.; Rocke, D. M.; Isseroff, R. R. Cellular versus Acellular Matrix Devices in the Treatment of Diabetic Foot Ulcers: Interim Results of a Comparative Efficacy Randomized Controlled Trial. J. Tissue Eng. Regen. Med. 2019, 13 (8), 1430–1437. 10.1002/term.2884.

(8) Imam, B.; Miller, W. C.; Finlayson, H. C.; Eng, J. J.; Jarus, T. Incidence of Lower Limb Amputation in Canada. Can. J. Public Heal. 2017, 108 (4), e374–e380. 10.17269/cjph.108.6093.

(9) Schaper, N. C.; Netten, J. J.; Apelqvist, J.; Bus, S. A.; Hinchliffe, R. J.; Lipsky, B. A. Practical Guidelines on the Prevention and Management of Diabetic Foot Disease (IWGDF 2019 Update). Diabetes. Metab. Res. Rev. 2020, 36 (S1), e3266. 10.1002/dmrr.3266.

(10) Santema, T. B.; Lenselink, E. A.; Balm, R.; Ubbink, D. T. Comparing the Meggitt-Wagner and the University of Texas Wound Classification Systems for Diabetic Foot Ulcers: Inter-Observer Analyses. Int. Wound J. 2016, 13 (6), 1137–1141. 10.1111/IWJ.12429.

(11) Fu, T.; Stupnitskaia, P.; Matoori, S. Next-Generation Diagnostic Wound Dressings for Diabetic Wounds. ACS Meas. Sci. 2022, 2, 377–384.

(12) McArdle, C.; Lagan, K. M.; McDowell, D. A. The PH of Wound Fluid in Diabetic Foot Ulcers -- the Way Forward in Detecting Clinical Infection? Curr. Diabetes Rev. 2014, 10 (3), 177–181.

(13) McArdle, C.; Lagan, K.; Spence, S.; McDowell, D. Diabetic Foot Ulcer Wound Fluid: The Effects of PH on DFU Bacteria and Infection. J. Foot Ankle Res. 2015, 8 (S1), A8. 10.1186/1757-1146-8-s1-a8.

(14) McArdle, C. D.; Lagan, K. M.; McDowell, D. A. Effects of PH on the Antibiotic Resistance of Bacteria Recovered from Diabetic Foot Ulcer Fluid An In Vitro Study. J. Am. Podiatr. Med. Assoc. 2018, 108 (1), 6–11. 10.7547/16-033.

(15) Percival, S. L.; McCarty, S.; Hunt, J. A.; Woods, E. J. The Effects of PH on Wound Healing, Biofilms, and Antimicrobial Efficacy. Wound Repair Regen. 2014, 22 (2), 174–186. 10.1111/WRR.12125.

(16) Mai, H.; Wang, Y.; Li, S.; Jia, R.; Li, S.; Peng, Q.; Xie, Y.; Hu, X.; Wu, S. A PH-Sensitive near-Infrared Fluorescent Probe with Alkaline PKa for Chronic Wound Monitoring in Diabetic Mice. Chem. Commun. 2019, 55 (51), 7374–7377. 10.1039/C9CC02289A.

(17) Tang, N.; Zhang, R.; Zheng, Y.; Wang, J.; Khatib, M.; Jiang, X.; Zhou, C.; Omar, R.; Saliba, W.; Wu, W.; Yuan, M.; Cui, D.; Haick, H. Highly Efficient Self-Healing Multifunctional Dressing with Antibacterial Activity for Sutureless Wound Closure and Infected Wound Monitoring. Adv. Mater. 2022, 34 (3), 2106842. 10.1002/ADMA.202106842.

(18) Schreml, S.; Meier, R. J.; Kirschbaum, M.; Kong, S. C.; Gehmert, S.; Felthaus, O.; Küchler, S.; Sharpe, J. R.; Wöltje, K.; Weiß, K. T.; Albert, M.; Seidl, U.; Schröder, J.; Morsczeck, C.; Prantl, L.; Duschl, C.; Pedersen, S. F.; Gosau, M.; Berneburg, M.; Wolfbeis, O. S.; Landthaler, M.; Babilas, P. Luminescent Dual Sensors Reveal Extracellular PH-Gradients and Hypoxia on Chronic Wounds That Disrupt Epidermal Repair. Theranostics 2014, 4 (7), 721–735. 10.7150/THNO.9052.

(19) Mordon, S.; Maunoury, V.; Devoisselle, J. M.; Abbas, Y.; Coustaud, D. Characterization of Tumorous and Normal Tissue Using a PH-Sensitive Fluorescence Indicator (5,6-Carboxyfluorescein) in Vivo. J. Photochem. Photobiol. B Biol. 1992, 13 (3–4), 307–314. 10.1016/1011-1344(92)85070-B.

(20) Matoori, S.; Bao, Y.; Schmidt, A.; Fischer, E. J.; Ochoa-Sanchez, R.; Tremblay, M.; Oliveira, M. M.; Rose, C. F.; Leroux, J.-C. An Investigation of PS-b-PEO Polymersomes for the Oral Treatment and Diagnosis of Hyperammonemia. Small 2019, 15 (50), e1902347. 10.1101/631630.

(21) Wang, S.; Peng, Y. Natural Zeolites as Effective Adsorbents in Water and Wastewater Treatment. Chem. Eng. J. 2010, 156 (1), 11–24. 10.1016/J.CEJ.2009.10.029.

(22) Albertson, T.; Derlet, R. W.; Foulke, G. E.; Minguillon, M.; Tharratt, S. Superiority of Activated Charcoal Alone Compared with Ipecac and Activated Charcoal in the Treatment of Acute Toxic Ingestions. Ann. Emerg. Med. 1989, 18 (1), 56–59. 10.1016/S0196-0644(89)80314-2.

(23) Augst, A. D.; Kong, H. J.; Mooney, D. J. Alginate Hydrogels as Biomaterials. Macromol. Biosci. 2006, 6 (8), 623–633. 10.1002/mabi.200600069.

(24) Matoori, S.; Veves, A.; Mooney, D. J. Advanced Bandages for Diabetic Wound Healing. Sci. Transl. Med. 2021, 13 (585), eabe4839.

(25) Nguyen, T. T.; Ding, D.; Wolter, W. R.; Pérez, R. L.; Champion, M. M.; Mahasenan, K. V.; Hesek, D.; Lee, M.; Schroeder, V. A.; Jones, J. I.; Lastochkin, E.; Rose, M. K.; Peterson, C. E.; Suckow, M. A.; Mobashery, S.; Chang, M. Validation of Matrix Metalloproteinase-9 (MMP-9) as a Novel Target for Treatment of Diabetic Foot Ulcers in Humans and Discovery of a Potent and Selective Small-Molecule MMP-9 Inhibitor That Accelerates Healing. J. Med. Chem. 2018, 61 (19), 8825–8837. 10.1021/acs.jmedchem.8b01005.

(26) Castleberry, S. A.; Almquist, B. D.; Li, W.; Reis, T.; Chow, J.; Mayner, S.; Hammond, P. T. Self-Assembled Wound Dressings Silence MMP-9 and Improve Diabetic Wound Healing In Vivo. Adv. Mater. 2016, 28 (9), 1809–1817. 10.1002/adma.201503565.

