## Supplementary Table and Figures for "Fluorescent pH-sensing bandage for point-of-care wound diagnostics"

**Corresponding author**

Simon Matoori

Université de Montréal

2940 Chemin de Polytechnique, Montreal, QC H3T 1J4

**Supplementary Table**

Supplementary Table S1. Size of microparticles in µm (n=3).

|  | **Activated charcoal MP** | **Zeolite MP** | **Silica MP** | **Carboxylate MP** | **BTA MP** |
| --- | --- | --- | --- | --- | --- |
| **d10** | 5.9 | 4.6 | 14.1 | 112.6 | 453.5 |
| **d50** | 24.0 | 14.3 | 49.3 | 158.7 | 660.1 |
| **d90** | 75.5 | 331.8 | 172.4 | 201.7 | 904.4 |

**Supplementary Figures**

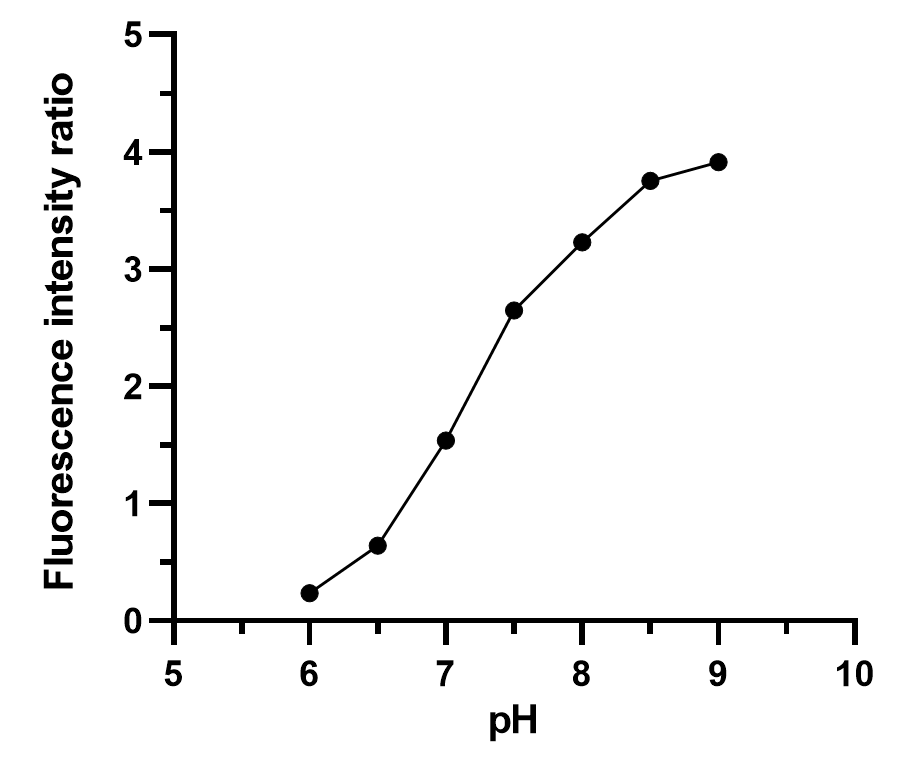

**Supplementary Figure S1.** pH sensitivity of pyranine. Fluorescence intensity ratio of pyranine at different pH values. All results as means ± SD (n = 3).

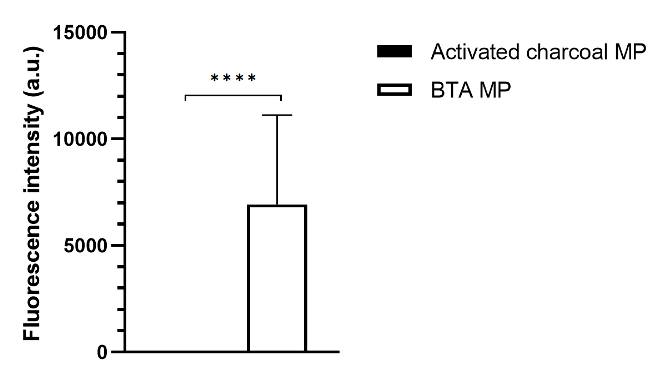

**Supplementary Figure S2.** Fluorescence intensity of pyranine-loaded activated charcoal MP and BTA microparticles. Pyranine fluorescence intensity at 413 nm (excitation wavelength) and 510 nm (emission wavelength). All results as means ± SD (n = 3). ****p < 0.0001.

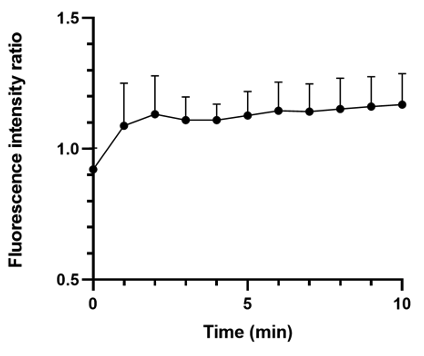

**Supplementary Figure S3.** Kinetics of pH sensing of pyranine-loaded BTA microparticles-encapsulating calcium alginate hydrogel. Pyranine fluorescence intensity ratio over time. Incubation temperature: 37C. All results as means ± SD (n = 3).
